# Prenatal thyroid hormones accelerate postnatal growth and telomere shortening in wild great tits

**DOI:** 10.1101/2021.12.22.473794

**Authors:** Bin-Yan Hsu, Nina Cossin-Sevrin, Antoine Stier, Suvi Ruuskanen

## Abstract

Early-life environment is known to affect later-life health and disease, which could be mediated by the early-life programming of telomere length, a key hallmark of ageing. According to the *fetal programming of telomere biology hypothesis*, variation in prenatal exposure to hormones is likely to influence telomere length. Yet the contribution of key metabolic hormones, *i.e*. thyroid hormones (THs), has been largely ignored. We recently showed that in contrast to predictions, exposure to elevated prenatal THs increased postnatal telomere length in wild collared flycatchers, but the generality of such effect, its underlying proximate mechanisms and consequences on survival have not been investigated. We therefore conducted a comprehensive study evaluating the impact of THs on potential drivers of telomere dynamics (growth, post-natal THs, mitochondria and oxidative stress), telomere length and medium-term survival using wild great tits as a model system. While prenatal THs did not significantly affect telomere length a week after hatching (*i.e*. day 7), they influenced postnatal telomere shortening (*i.e*. shorter telomeres at day 14 and the following winter) but not apparent survival. Circulating THs, mitochondrial density or oxidative stress biomarkers were not significantly influenced, whereas TH-supplemented group showed accelerated growth, which may explain the observed delayed effect on telomeres. We discuss several alternative hypotheses that may explain the contrast with our previous findings in flycatchers. Given that shorter telomeres in early life tend to be carried until adulthood and are often associated with decreased survival prospects, the effects of prenatal THs on telomeres may have long-lasting effects on senescence.

## Introduction

Early-life environment has been repeatedly observed to affect adult health and survival prospects in human and non-human vertebrates (Barnes and Ozanne, 2011; Godfrey and Barker, 2001; Metcalfe and Monaghan, 2001). While the mechanisms underlying such delayed effects remained somewhat elusive (Barnes and Ozanne, 2011), the early-life programming of telomere length (i.e. the protective end caps of chromosomes) has emerged as a key candidate (Entringer et al., 2018). Telomere length is considered as a hallmark of ageing (López-Otín et al., 2013) since telomeres shorten with age, and shorter telomeres are often predictive of lower survival or lifespan in both epidemiological and experimental studies (Arbeev et al., 2020; Heidinger et al., 2012; Muñoz-Lorente et al., 2019; Wilbourn et al., 2018). The prenatal hormonal environment, such as exposure to elevated glucocorticoid levels, has been coined as an important factor influencing early-life telomere length and its associated long-term outcomes (Criscuolo et al., 2020; Haussmann et al., 2012; Marchetto et al., 2016; Parolini et al., 2019). While there has been a considerable interest in prenatal glucocorticoids (Haussmann et al. 2012; Tissier, Williams, and Criscuolo 2014; Noguera, da Silva, and Velando 2020) and to a lesser extent androgens (Parolini et al., 2019; Tissier et al., 2014) in the context of the *‘fetal programming of telomere biology’* hypothesis (Entringer et al., 2018), the potential impact of prenatal thyroid hormones has been mostly ignored so far (Stier et al. 2020).

Thyroid hormones (THs) are key coordinators of development and metabolism (McNabb, 2007), which are transferred from mothers to offspring (Ruuskanen and Hsu, 2018). Variation in exposure to prenatal thyroid hormones (T3, triiodothyronine, and T4, thyroxine) could influence telomere length via several mutually non-exclusive proximate pathways: (i) Prenatal THs can stimulate growth (yet results are inconsistent across studies and species, (Hsu et al., 2020, 2019a, 2017; Medici et al., 2013; Ruuskanen et al., 2016a; Sarraude et al., 2020a; Sarraude et al., 2020b; Vrijkotte et al., 2017; Zhang et al., 2019), which can directly contribute to telomere attrition through increasing cellular division (Monaghan and Ozanne, 2018; Stier et al., 2020), or indirectly accelerate telomere shortening through increasing oxidative stress (Monaghan and Ozanne, 2018; Reichert and Stier, 2017; Smith et al., 2016). (ii) Elevated TH levels are often associated with higher metabolic rates (Liu et al., 2006; Mullur et al., 2014; Welcker et al., 2013) and stimulate mitochondrial aerobic metabolism (Cioffi et al., 2013), both of which can potentially increase reactive oxygen species (ROS) and oxidative damage (Stier, Massemin, and Criscuolo 2014), accelerating telomere shortening (Reichert and Stier, 2017). It was recently shown that exposure to elevated prenatal TH levels can lead to a sex-specific increase in metabolic rate and circulating TH levels shortly after hatching (rock pigeons *Columba livia*, Hsu et al. 2017, but see Ruuskanen et al. 2016), which suggests that prenatal hormones may program postnatal metabolism and TH-axis function. (iii) The ‘metabolic telomere attrition hypothesis’ (Casagrande and Hau, 2019) postulates that telomere shortening might be adaptive by amplifying the cellular signaling of energy debt to re-direct critical resources to immediately important processes. In this scenario, we would expect accentuated shortening when catabolism is favored over anabolism via mTOR inhibition (i.e. mechanistic target of rapamycin, a key regulator of cellular homeostasis) leading to a decrease in telomere maintenance processes and an active shortening through exonuclease action (Casagrande and Hau, 2019). Since THs can have both anabolic and catabolic actions (Mullur et al., 2014), predictions can be made in both directions between prenatal THs and telomere length.

From an evolutionary perspective, increased offspring metabolism or growth, diverts resources from somatic maintenance if resources are limited. This can accelerate damage to biomolecules and/or decrease repair/maintenance processes, and therefore accentuate telomere shortening. Therefore, prenatal TH levels would be expected to vary in relation to predicted environmental conditions (as observed in terms of temperature and laying order, e.g. Ruuskanen, Groothuis, et al. 2016; Hsu, Verhagen, et al. 2019) to optimize this trade-off.

In contrast to most predictions relating prenatal THs to telomere length (see above), we recently reported that prenatal exposure to experimentally elevated THs increased telomere length in nestlings of a wild passerine, the collared flycatcher (*Ficedula albicollis*, Stier et al. 2020). To better understand the potential generality of this surprising finding as well as assess underlying mechanisms and potential carry-over effects of variation in prenatal THs on later life-stages and survival, we conducted a more detailed study in another passerine species, the great tit *(Parus major*). The aim of this study was to comprehensively investigate the influence of prenatal THs on growth, oxidative stress, plasma THs, mitochondrial density (i.e. mitochondrial DNA copy number) and telomere dynamics as well as survival via an experimental manipulation of prenatal THs in a wild population. We monitored offspring multiple times during development and as juveniles a few months after fledging. Based on the majority of prior literature, we would predict that elevated prenatal THs could lead to faster growth, increased plasma THs, oxidative stress and mitochondrial density (higher density can lead to higher ROS production), ultimately accelerating telomere shortening. Alternatively, if our previous finding in the collared flycatcher reflected a general pattern, we would predict that despite accelerating growth, elevated prenatal THs could increase telomere length. We also predict that elevated prenatal TH could increase post-fledging survival of the juveniles (*e.g*. both due to accelerated postnatal growth and potential beneficial effects for thermoregulation under low autumn-winter temperatures). However, longer-term survival, that we were not able to evaluate here, could be decreased for example due to shorter telomeres.

## Methods

The experiment and all methods we used were in accordance with all relevant guidelines and regulations and have been approved by the Animal Experiment Board of the Administrative Agency of South Finland (ESAVI/2902/2018) and the Environmental Center of Southwestern Finland (license number VARELY549/2018). The experiment was conducted in 2018 in a nest box population (314 nest-boxes distributed over seven forest plots) on the island of Ruissalo in southwestern Finland (60° 25’ N, 22° 10’ E).

### Field experiment

The nest boxes were monitored with five-day intervals to track egg laying. Yolk T3 and T4 (i.e. a combined injection of the two hormones) levels were elevated in half of the nests (n = 21 TH nests and n = 21 control nests) by injection into the egg, following methods in Ruuskanen, Darras, et al. (2016). Control nests were injected with the vehicle (0.9% NaCl) only. The TH content (±SD) in great tit eggs in the population is 0.053 (±0.02) ng/yolk for T3 and 0.46 (± 0.16 ng/yolk) for T4 (Ruuskanen et al. unpubl data). We aimed to raise the amount of yolk TH by 2SD via injection into the egg yolk, a dose that has been recommended in relation to the natural hormone range of the study species (Podmokla et al., 2018). This corresponded to target doses of 0.041 ng/yolk for T3 and 0.325 ng/yolk for T4. To make sure injections would be performed on unincubated eggs mimicking maternal hormone levels (great tits can start incubation before the clutch is complete), injections were conducted on the day the 5^th^ egg was laid to all eggs in the clutch. Thereafter, injections were conducted each day for the newly laid egg.

Hatching date and success was monitored by visiting nests daily starting before the estimated hatching day. Nestling body mass (~0.01g) and tarsus (~0.5 mm) were measured on day 2 (mass only), 7 and 14 post-hatching. All nestling measurements were conducted between 8 am and 2 pm. After measurements on day 2, nestlings were nail-coded, the brood was split and nestlings were cross-fostered with same-age nestlings in another nest of the opposite hormone treatment. This “split-brood design” allows chicks from TH and control-injected eggs to be raised under the same postnatal environment and is in theory more sensitive to detect effects that may otherwise require a much larger number of nests in an experiment of “between-brood” design (e.g. Tschirren et al. 2005). Preferentially, half of the brood was swapped between the nests with opposite treatments whenever possible, but we also split a brood for cross-fostering when we did not have an equal number of nests with opposite treatments on a given day. The proportion of TH-nestlings (i.e. number of TH-nestlings divided by the brood size) in each nest after cross-fostering was therefore recorded. All nestlings, cross-fostered or not, were included in the statistical analyses. There was no clear bias on the number of nestlings that were cross-fostered or stayed in the original nests with respect to the hormone treatment (CO-nestlings: 58 stayed and 41 moved, TH-nestlings: 68 stayed and 51 moved, χ^2^ = 0.006, p = 0.939). On day 7 a blood sample (ca 40μl) was taken and snap-frozen in liquid nitrogen for molecular analyses (telomere length, mitochondrial density and molecular sexing). On day 14 a small blood sample (10-15μl) was snap-frozen in liquid nitrogen for oxidative stress biomarker analyses and stored at −80°C, while the rest of the sample (ca 40 μl) was kept on ice, and centrifuged. RBCs (15 μl) were used for mitochondria and telomere measurements as above and plasma (15 μl) for TH analyses. After day 7 measurements half of the nest were subject to a temperature manipulation (nest temperature increased for *ca*. 2°C during day 7 – day 14) for the purposes of another study. To avoid potential confounding effects, for data after day 7 and in juveniles, we only included the individuals from the nests which were not temperature-manipulated (N = 20 nests, 111 nestlings on day 7 and 99 nestlings on day 14). Yet, for day 2 and day 7 measurements (prior to the temperature manipulation), we prefer to include all data (N = 42 nests, in total 224 nestlings, 218 nestlings on d2 and 221 nestlings on d7) to make use of the larger sample size.

To study long-term effects of prenatal thyroid hormones and offspring apparent post-fledging survival, we recaptured great tits during the following autumn-winter using mist-netting. Seven feeding stations were mounted in the study plots in August (generally any given nest box had a station within 200m), and nets were positioned close to the feeding stations. The distances between adjacent feeding stations were ca. 500 m, and most birds were captured at a different feeding station than their natal forest plot. In addition, 5 birds were captured at a constant effort site > 3 km from the study plots, suggesting that all stations were potentially accessible to all birds. Circa 20 kg of peeled sunflower seeds and 2 kg of fat were provided at each station, checked and filled bi-weekly, and consumption noted. Only in a very few cases the feeders were completely empty. Mist-netting was conducted at each feeding station for 3 hours on three different days in September-October and similarly again in February, summing up to a total 126 hours of mist-netting. The time of day (morning/day/afternoon) was rotated for each site. Nets were checked every 30 minutes and mass (~0.01g), wing (~0.5 mm) were recorded for each bird. A small blood sample (40-60μl) was taken, kept on ice, centrifuged within 8 hours, and RBCs frozen at −80°C for telomere and mitochondria density analyses.

### Thyroid hormones and oxidative stress assays

Plasma thyroid hormones, the biologically active form triiodothyronine (T3), and a precursor thyroxine (T4) (expressed as ng/mL) were measured from 14-day-old nestlings with nano-LC-MS/MS following Ruuskanen et al. (2019; 2018). Due to practical constraints (time, effort and finances), we randomly selected one to four nestling per nest, n = 13 TH and n = 11 control nestlings (from 18 nests of origin and 11 nests of rearing) for analysis.

Total glutathione (tGSH), the most abundant intra-cellular antioxidant, and the ratio between reduced and oxidized gluthatione (GSH:GSSG, an indicator of oxidative challenge) were measured from whole-blood samples (12μl of 0.9% NaCl-diluted whole blood) with the ThioStar® Glutathione Fluorescent Detection Kit (K005-FI, Arbor Assays, USA) (Sarraude et al., 2020a). As a measure of oxidative damage, we assessed blood lipid peroxidation (malonaldehyde, MDA, 50μl of 0.9% NaCl-diluted whole blood) using the TBARS-assay following Espin et al. (2017). Both measurements had CV% < 10 and are expressed per mg of protein (measured via BCA protein assay, ThermoFisher Scientific), following Espin et al. 2017. We aimed to measure biomarkers of oxidative stress from 2 control or 2 TH-nestlings per nest, yet sometimes fewer nestlings were available and some samples failed in the laboratory analysis, yielding the final sample size as n = 33 TH and 26 control nestlings from 18 rearing nests.

### qPCR assays for relative telomere length, mtDNA copy number and molecular sexing

We extracted DNA from blood cells using a standard salt extraction alcohol precipitation method (Aljanabi and Martinez, 1997). Extracted DNA was diluted in elution buffer BE for DNA preservation. DNA concentration and quality (260/280 > 1.80 and 260/230 > 2.00) were checked with a ND-1000-Spectrophotometer (NanoDrop Technologies, Wilmington, USA). DNA integrity was verified in 24 samples chosen randomly using gel electrophoresis (50 ng of DNA, 0.8 % agarose gel at 100 mV for 60 min) and DNA staining with Midori Green. Each sample was then diluted to a concentration of 1.2 ng.μl^-1^ for subsequent qPCR analysis.

Relative telomere length (*rTL*) and mitochondrial DNA copy number (*mtDNAcn*, an index of mitochondrial density) were quantified using qPCR. This technique estimates relative telomere length by determining the ratio (T/S) of telomere repeat copy number (T) to a single copy gene (SCG), and the relative mtDNAcn as the ratio between one mitochondrial gene and the same single copy gene. Here, we used RAG1 as a SCG (verified as single copy using a BLAST analysis on the great tit genome) and cytochrome oxidase subunit 2 (COI2) as a mitochondrial gene (verified as non-duplicated in the nuclear genome using a BLAST analysis). Forward and reverse telomere primers were 5’-CGGTTTGTTTGGGTTTGGGTTTGGGTTTGGGTTTGGGTT-3’ (Tel-1b) and 5’-GGCTTGCCTTACCCTTACCCTTACCCTTACCCTTACCCT-3’ (Tel-2b) respectively. Forward and reverse RAG1 primers were 5’-TCGGCTAAACAGAGGTGTAAA-3’ and 5’-CAGCTTGGTGCTGAGATGTAT-3’, respectively. Forward and reverse COI2 primers were 5’-CAAAGATATCGGCACCCTCTAC-3’; 5’-GCCTAGTTCTGCACGGATAAG-3’, respectively. For the qPCR assays, the reactions were performed on a 384-QuantStudio™ 12K Flex Real-Time PCR System (Thermo Fisher), in a total volume of 12μL including 6ng of DNA, primers at a final concentration of 300nM and 6μL of SensiFAST™ SYBR lo-ROX (Bioline). Telomere, RAG1 and COI2 reactions were performed in triplicates on the same plates (10 plates in total); the qPCR conditions were: 3min at 95°C, followed by 35 cycles of 10 s at 95°C, 15 s at 58°C and 10s at 72°C. A DNA sample being a pool of DNA from 10 adult individuals was used as a reference sample and was included in triplicate on every plate. The efficiency of each amplicon was estimated from a standard curve of the reference sample ranging from 1.5 to 24ng. The mean reaction efficiencies were 109.1 ± 1.8% for telomere, 102.2 ± 1.6% for RAG1, 96.3 ± 1.1% for COI2. The relative telomere length and mtDNAcn of each sample were calculated as (1+*Ef*_Tel or COI2_)^ΔCq Tel or COI2^/(1+*Ef*_RAG1_)^ΔCqRAG1^; *Ef* being the amplicon efficiency, and ΔCq the difference in Cq-values between the reference sample and the focal sample. Intra-plate technical repeatabilities of telomere and *mtDNAcn* based on triplicates were 0.87 (95% C.I. [0.85-0.89]) and 0.96 (95% C.I. [0.95-0.97]) respectively. Inter-plate technical repeatabilities of telomere and *mtDNAcn* based on one repeated plate were 0.98 (95% C.I. [0.97-0.99]) and 0.77 (95% C.I. [0.59-0.88]) respectively.

The use of *mtDNAcn* as an index of mitochondrial density has been questioned in human (Larsen et al., 2012), but we have previously shown good correlations between mtDNAcn and mitochondrial respiration rates in pied flycatcher (Stier et al. 2019). Great tits have quite peculiar telomeres, characterized notably by some ultra-long telomeres that do not seem to shorten with age in adults (Atema et al., 2019). Since qPCR only provides an estimate of overall telomere length, it has been suggested it could be suboptimal for this study species. Yet, relative telomere length (*i.e*. measured using qPCR) in this species has been shown to shorten during the nestling stage (Stier et al., 2016, 2015), to respond to environmental factors (*e.g*. hatching asynchrony: (Salmón et al., 2016; Stier et al., 2016, 2015) and to predict adult survival (Salmón et al., 2017). Within-individual repeatability of telomere length has recently been suggested to be an important factor to evaluate the pertinence of telomere length data in a given study/species (Kärkkäinen et al., n.d.), and the biological repeatability in our dataset was *R* = 0.66 [0.55-0.79], which is above the average reported by qPCR studies (*i.e. R* = 0.47), and within the upper range of what has been reported for great tits (Kärkkäinen et al. 2021)

Birds were molecularly sexed using a qPCR approach adapted from(Chang et al., 2008; Ellegren and Fridolfsson, 1997). Forward and reverse sexing primers were 5’-CACTACAGGGAAAACTGTAC-3’ (2987F) and 5’-CCCCTTCAGGTTCTTTAAAA-3’ (3112R), respectively. qPCR reactions were performed in a total volume of 12μL including 6ng of DNA, primers at a final concentration of 800nM and 6μL of SensiFAST™ SYBR® Lo-ROX Kit (Bioline). qPCR conditions were: 3 min at 95°C, followed by 40 cycles of 45 s at 95°C, 60 s at 52°C and 60s at 72°C, then followed by a melting curve analysis (95°C 60s, 45°C 50s, increase to 95°C at 0.1°C/s, 95°C 30s). Samples were run in duplicates in a single plate and 6 adults of known sex were included as positive controls. Sex was determined by looking at the dissociation curve, with two peaks indicating the presence of a Z and W chromosome (female), and one peak indicating the presence of only the Z chromosomes (male).

### Statistical analyses

We ran several linear or generalized linear mixed models (LMMs/GLMMs) for both nestling and juvenile data. In order to account for genetic effects and the effects of growing environment other than prenatal hormones, we included the IDs of both the nest of origin (i.e. where a nestling hatched) and the nest of rearing (i.e. where a nestling grew up) as random intercepts in all suitable models. For physiological measurements, assay batch ID was included either as random intercept or fixed factors (depending on the number of levels, see Tables for details). All initial models included the prenatal hormone treatment (TH vs. CO), date (continuous variable), cross-fostering (yes/no), brood size (continuous variable), proportion of TH-nestlings (continuous variable) in a nest, and their interactions with hormone treatment as fixed factors. Except for hormone treatment, other variables were initially included because of their potential confounding effects on nestling growth and development but only retained in the final models when p < 0.15 in order not to miss potentially significant factors. Nevertheless, in the final models, significance level was still set at 0.05. In models of nestling growth (i.e. body mass and tarsus length), the ID of each individual was included as a random intercept.

To analyze nestling growth, we separated the data into the early (i.e. from day 2 to day 7) and late (i.e. from day 7 to day 14) phases. Because of the start of heating treatment at day 7, this separation allows us to make use of the larger sample size during the early phase and focus on the nestlings from non-heated nests during the late phase. In these models, age (as a continuous variable) and hormone × interaction were included as additional fixed factors, and individual ID as a random intercept. In order to account for the variation in growth rate between nests, age (continuous variable) was also treated as a random slope to interact with both the original and rearing nests. For tarsus length, we additionally controlled for the measurer ID as a fixed factor. Because molecular sexing was only conducted on nestlings having DNA extracted for mtDNAcn and rTL, not all nestlings included in the growth analyses were sexed. We therefore did not include sex in order to make use of the full data set. Repeating our models with only the sexed nestlings still gave qualitatively the same results and did not show sex-specific effects of prenatal THs on growth. For the juveniles captured during autumn/winter we ran LMMs with hormone treatment, date of capture, and sex (because of clear sexual size dimorphism at this age) as fixed factors.

For all models of physiological measurements fixed factors included prenatal hormone treatment, nestling sex and body mass. If not showing any trends (i.e. p<0.15), sex and body mass were then removed from the final models, given the smaller sample sizes for the physiological measurements. Natural log transformation was conducted on highly-skewed data: mtDNAcn and three oxidative stress biomarkers to ensure normal residual distributions. rTL and plasma THs were untransformed as residuals qualified the assumption of normality.

Because both mtDNAcn and rTL are ratios to a single copy gene, we also z-transformed the data to allow across-study comparison (Verhulst, 2020). The ages of all juveniles were pooled as a single age category “autumn”, and age was thus included as a three-level categorical variable (day 7, day 14, and autumn) in order to estimate the changes of mtDNAcn and rTL between each time point.

We used GLMMs to estimate the influence of prenatal thyroid hormones on hatching success, fledging success (i.e. pre-fledging survival), and post-fledging survival. We modelled the outcome of each egg (i.e. hatched or not) or nestling (i.e. survived or not) using a logit link function and specifying a binomial residual distribution. For hatching success, two nests in which eggs never hatched (i.e. cause of failure likely unrelated to our treatment) were excluded, giving a final sample size of 354 eggs from 42 nests. The nest ID was treated as a random intercept, and prenatal hormone treatment and laying date of each nest as fixed factors. For fledging success, we only included the 131 nestlings (reared in 27 nests after cross-fostering) that successfully hatched and were subsequently reared in the nests that were not subject to temperature manipulation in the analysis. Among the 131 nestlings, 99 successfully fledged and were included in the analysis of post-fledging survival. According to the nestling ringing records since 1999, only 8 out of >6200 birds ringed on the Ruissalo island were recaptured outside the island, suggesting a very low dispersal rate. The individuals that were never recaptured in the autumn were therefore presumed dead. The ID of rearing and original nests was treated as a random intercept and prenatal hormone treatment as the fixed factor. Laying date, brood size, and cross-fostering, as well as their interactions with the hormone treatment, were initially included but only retained in the final model when p < 0.15.

In all statistical models described above, hormone-related two-way interactions were of interest and therefore included. Following the suggestion by (Schielzeth, 2010), input variables (except for the technical variables) were mean-centered before model fitting. Significant interactions were further examined by post-hoc interaction analysis using the R package *emmeans* (Lenth 2021).

All LMMs and GLMMs were conducted in the R environment 3.6.1, using the package *lme4* (Bates et al., 2015) and *lmerTest* (Kuznetsova et al., 2017). P values were determined using the Kenward-Roger method to approximate the denominator degrees of freedom (R package *pbkrtest*, (Halekoh and Hojsgaard, 2014), implemented within *lmerTest*) for LMMs and by Laplace approximation for GLMMs. Model assumptions of normality and homoscedasticity were diagnosed by visual inspection on simulated residuals using the R package *DHARMa* (Hartig 2021). Clear violation was only observed in the models for plasma T3 in which residuals were heteroscedastic. Nevertheless, given the fact that the model of T3 did not detect any significant effect and the general robustness of LMMs (Schielzeth et al., 2020), this did not influence our conclusion.

## Results

Early body mass growth (between day 2 and day 7 post-hatching) did not differ between TH-supplemented and control groups (Table 1a, Fig 1a), but individuals from TH-supplemented eggs grew faster at the later nestling stage, i.e. between day 7 and day 14 post-hatching (Hormone × age interaction, F_1,20.18_ = 5.449, p = 0.030, Table 1b, Fig 1b; body mass gain (mean±SE) in TH-nestlings 0.899 ± 0.042 g/day, CO-nestlings 0.786±0.043 g/day). Similarly, individuals from TH-supplemented eggs expressed marginally faster tarsus growth rate between day 7 and day 14 post-hatching (F_1,31.06_ = 3.737, p=0.062, Table 1c, TH: 0.567±0.033 mm/day, CO: 0.476±0.034 mm/day, Fig 1c). Cross-fostering was retained in the final model of early body mass growth and suggested that fostered chicks had a lower growth rate (−0.139±0.069 g/day) than non-fostered chicks (p = 0.046, Table 1a). When analyzing the body mass and tarsus separately for each age, body mass was slightly smaller on day 2 and day 7 in the TH-supplemented group compared to control, yet there was no statistically significant difference between the groups at any age (Fig 1, Table S1). At day 7, fostered nestlings tended to have a lower body mass by 0.347 g (SE = 0.179, p=0.053) on average, and had significantly shorter tarsi by 0.33 mm (SE = 0.146, p=0.025), yet such differences disappeared by day 14 (Table S1). In juveniles, there was no difference in body mass or wing length between the treatment groups (Body mass: F_1,9.79_ = 0.003, p = 0.959; wing length: F_1,10.34_ = 0.011, p = 0.919).

**Fig. 1.**
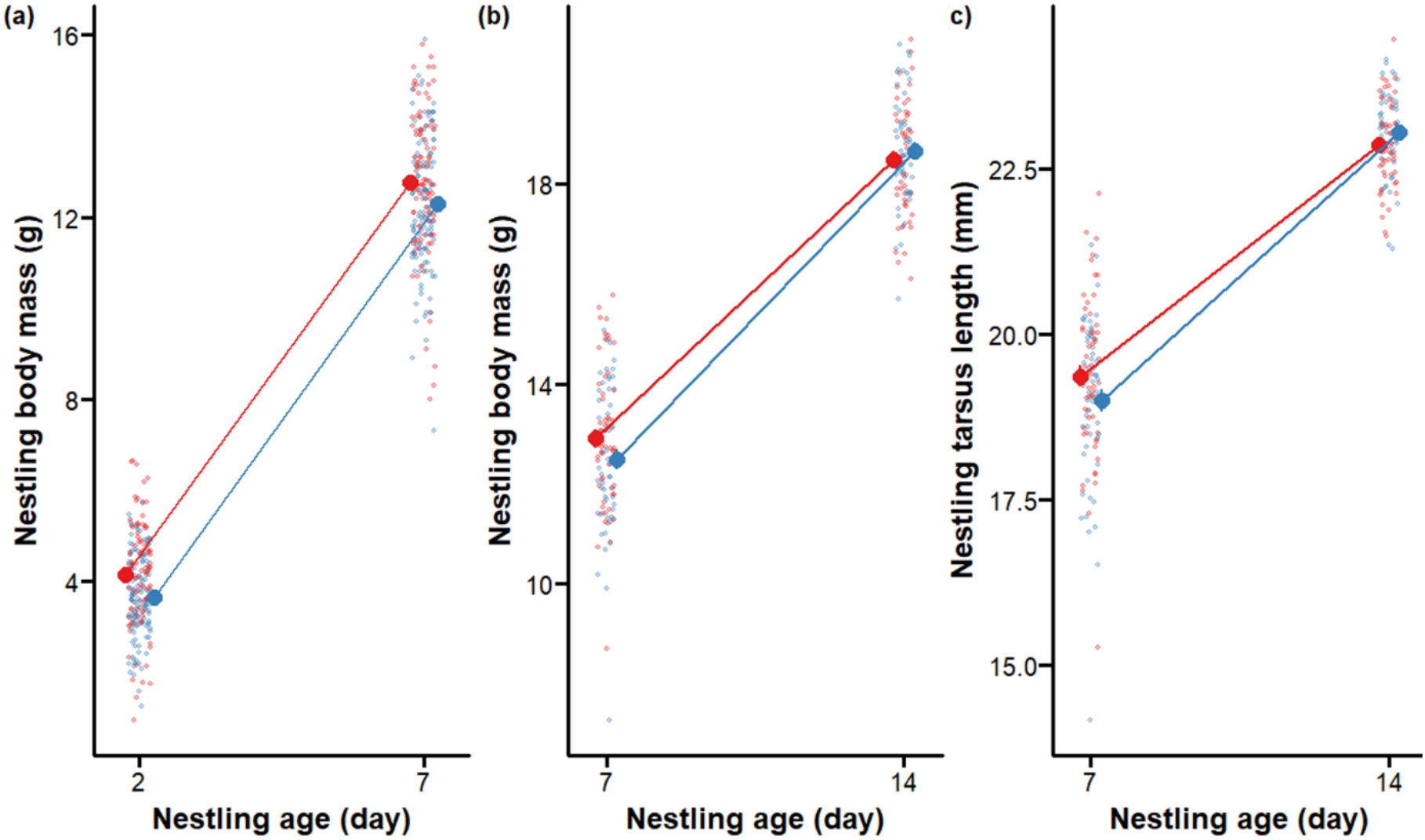
Nestling growth in prenatally TH-supplemented (blue dots) and control (red dots) groups. (a) body mass growth during early nestling stage (day 2 to day 7 post-hatching) (b) body mass growth late nestling stage (day 7 to day 14 post-hatching), (c) structural size (tarsus) growth during late nestling stages. Means±SE (large dots) and scatter of the raw data (small, semi-transparent dots) are shown. See sample sizes in the text.

**Table 1.**
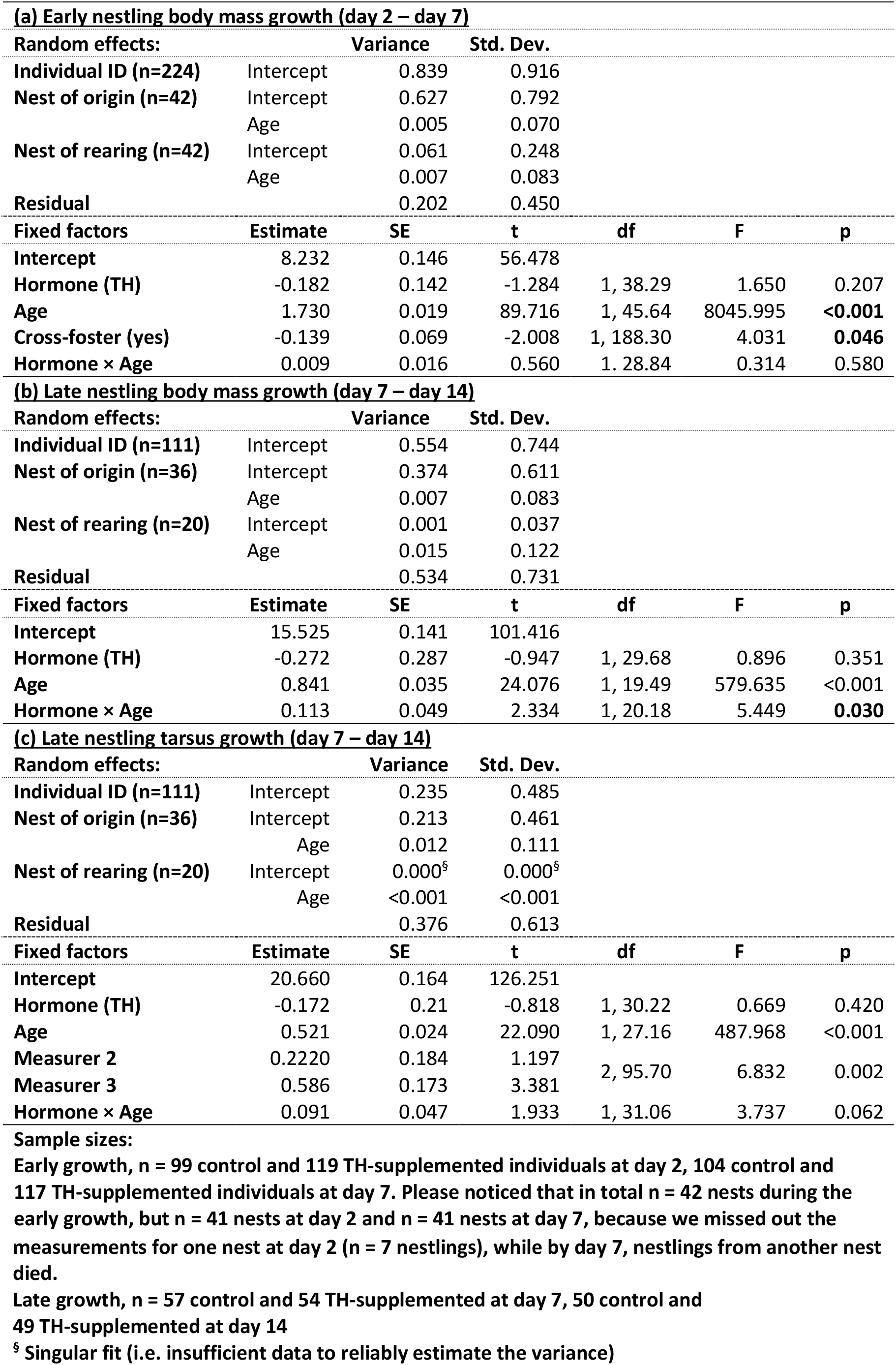
Linear mixed models of early and late nestling growth in response to prenatal TH-supplementation.

Biomarkers of intracellular oxidative status (total GSH, GSH:GSSG ratio) and damage to lipids 14 days post-hatching did not clearly differ among the treatment groups (marginal means±SE: total GSH μmol/mg protein: TH-supplemented, 0.253±0.022, control, 0.249±0.023; GSH:GSSG ratio: TH-supplemented, 0.045±0.008, control, 0.053±0.010; MDA nmol/mg protein: TH-supplemented, 0.038±0.005, control, 0.037±0.005; Table S2). Plasma T3 and T4 also did not differ among prenatally TH-supplemented and control groups 14 days post hatching (marginal means±SE pmol/ml T3: TH-supplemented, 1.21±0.17, control, 1.23±0.19; T4: TH-supplemented, 7.35±0.946, control, 7.39±1.02; Table S3).

Mitochondrial density decreased with age, but there were no clear differences between prenatally TH-supplemented and control groups at any age (d7, d14 or juveniles, Fig 2a, Table 2). Post-hoc analyses indicated significant decrease between each age category (all Tukey-adjusted p < 0.001), regardless of the hormone treatment.

**Fig 2.**
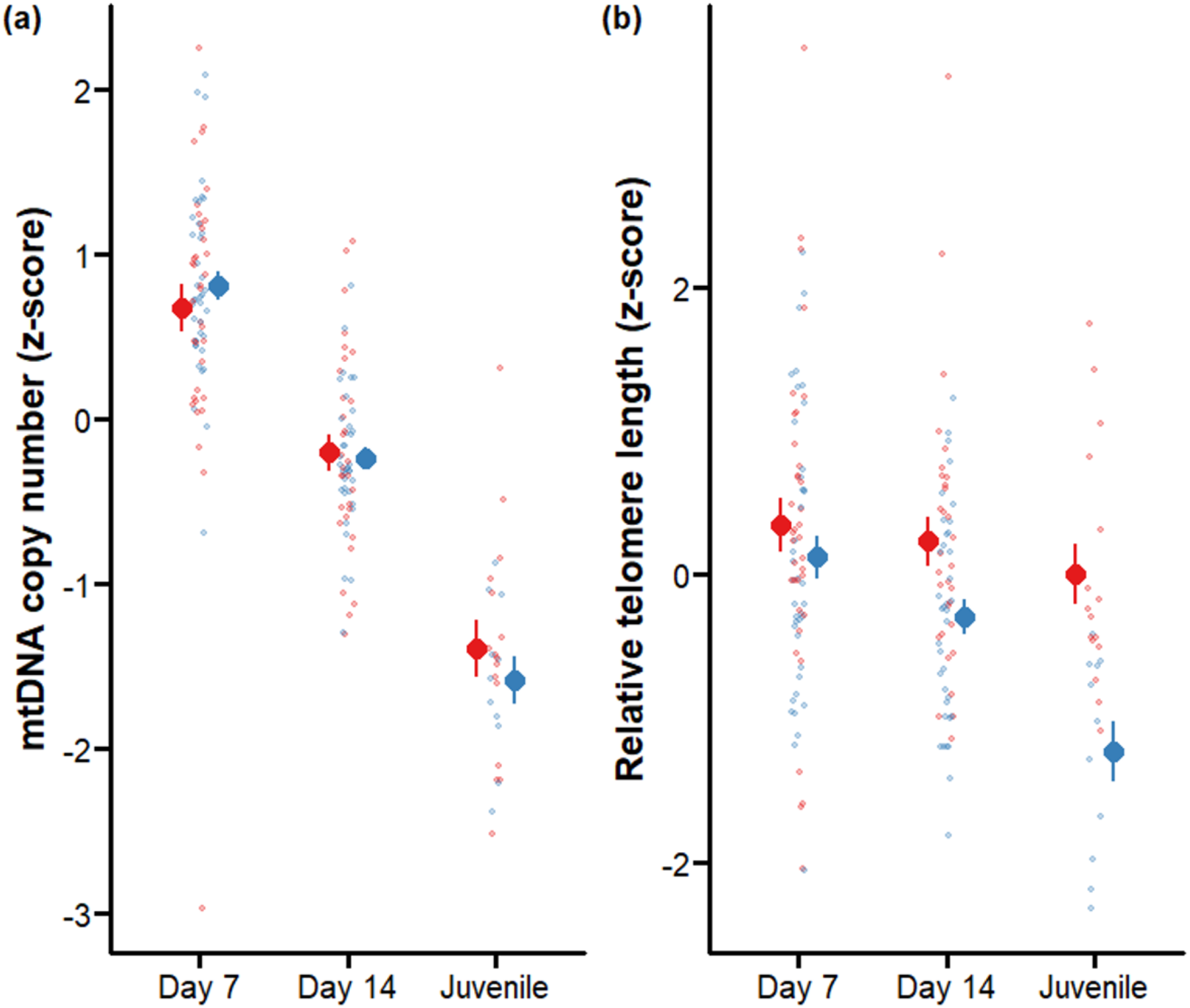
(a) Relative mitochondrial density (z-scale) and (b) relative telomere length in prenatally TH-supplemented (blue dots) and control groups (red dots) at different developmental stages (day 7 and 14 post-hatching and as juveniles, ca. 3 months of age). Means±SE (large dots) and scatter of the raw data (small, semi-transparent dots) are shown. See sample sizes in the text.

**Table 2.** Linear mixed models of mitochondrial density and telomere length in blood cells growth in response to prenatal TH-supplementation. In these models, age was as a categorical factor with three levels. For the purpose of mean-centering, two dummy variables (“Age day 7” and “Age day 14”) were created to represent the respective contrast against the juvenile age.

| **(a) Mitochondria density** |  |  |  |  |  |  |
| --- | --- | --- | --- | --- | --- | --- |
| **Random effects:** |  | **Variance** | **Std. Dev.** |  |  |  |
| **Individual ID (n=78)** | Intercept | 0.084 | 0.290 |  |  |  |
| **Nest of origin (n=35)** | Intercept | 0.041 | 0.203 |  |  |  |
| **Nest of rearing (n=19)** | Intercept | 0.000<sup>§</sup> | 0.000<sup>§</sup> |  |  |  |
| **DNA batch (n=8)** | Intercept | 0.017 | 0.129 |  |  |  |
| **Residual** |  | 0.244 | 0.494 |  |  |  |
| **Fixed factors** | **Estimate** | **SE** | **t** | **df** | **F** | **p** |
| Intercept | -0.023 | 0.079 | -0.295 |  |  |  |
| Hormone (TH) | 0.021 | 0.135 | 0.155 | 1, 24.00 | 0.024 | 0.878 |
| Sex | 0.200 | 0.121 | 1.651 | 1, 73.16 | 2.723 | 0.103 |
| Age day 7 | 2.202 | 0.121 | 18.134 | 1, 111.27 | 328.825 | **<0.001** |
| Age day 14 | 1.225 | 0.123 | 9.968 | 1, 111.12 | 99.357 | **<0.001** |
| Hormone × Age day 7 | 0.352 | 0.240 | 1.464 | 1, 111.78 | 2.142 | 0.146 |
| Hormone × Age day 14 | 0.197 | 0.243 | 0.807 | 1, 111.70 | 0.652 | 0.421 |
| **(b) relative telomere length** |  |  |  |  |  |  |
| **Random effects:** |  | **Variance** | **Std. Dev.** |  |  |  |
| **Individual ID (n=77)** | Intercept | 0.093 | 0.305 |  |  |  |
| **Nest of origin (n=35)** | Intercept | 0.000<sup>§</sup> | 0.000<sup>§</sup> |  |  |  |
| **Nest of rearing (n=19)** | Intercept | 0.026 | 0.162 |  |  |  |
| **DNA batch (n=8)** | Intercept | 0.394 | 0.628 |  |  |  |
| **Residual** |  | 0.513 | 0.716 |  |  |  |
| **Fixed factors** | **Estimate** | **SE** | **t** | **df** | **F** | **p** |
| Intercept | -0.085 | 0.236 | -0.361 |  |  |  |
| Hormone (TH) | -0.635 | 0.153 | -4.150 | 1, 16.06 | 17.223 | **<0.001** |
| Sex | -0.268 | 0.154 | -1.742 | 1, 65.94 | 3.035 | 0.086 |
| Age day 7 | 0.904 | 0.174 | 5.197 | 1, 117.27 | 27.008 | **<0.001** |
| Age day 14 | 0.654 | 0.175 | 3.728 | 1, 115.66 | 13.895 | **<0.001** |
| Brood size at day 2 | 0.213 | 0.094 | 2.271 | 1, 10.18 | 5.157 | **0.046** |
| Hormone × Age day 7 | 1.039 | 0.343 | 3.027 | 1, 120.32 | 9.165 | **0.003** |
| Hormone × Age day 14 | 0.741 | 0.347 | 2.137 | 1, 119.21 | 4.567 | **0.035** |
| Hormone × Brood size | -0.255 | 0.151 | -1.681 | 1, 32.83 | 2.827 | 0.102 |

Sample sizes: n = 37 control and 41 TH-supplemented individuals at day 7; 33 control and 38 TH-supplemented individuals at day 14; 16 control and 13 TH-treatment recaptured juveniles
<sup>§</sup> Singular fit (i.e. insufficient data to reliably estimate the variance)

Telomere length decreased with age (Table 2, Fig 2b), but the pattern was different between TH-supplemented and control group (Table 2, Fig 2b): Early in the nestling phase the difference between the prenatally TH-supplemented and control group approached significance (d7, Tukey post-hoc test: TH vs control: t_46.2_ = 1.804, p = 0.078). On day 14 post-hatching, telomeres were significantly shorter (13%) in the TH group (Tukey post-hoc test, TH vs Control: t_46.8_ = 3.166, p = 0.003), and this difference was accentuated in juveniles, offspring from TH supplemented eggs having telomeres being ca. 33% shorter than offspring from control group (Tukey post-hoc test, TH vs Control: t_109.8_ = 4.311, p < 0.0001). Further, we added the percentage of body mass gain from day 7 to day 14 in the model to examine whether the effect of prenatal TH supplementation was via its effect on nestling growth, but this did not qualitatively change the results. The percentage of body mass gain did not show a clear effect on telomere length (estimate±SE = −0.575±0.580, F_1,64.54_=0.981, p=0.326). Similarly, the three oxidative stress biomarkers also did not explain the difference in telomere shortening (all p’s > 0.31).

Hatching success did not clearly differ among prenatally TH supplemented and control individuals (TH 67.6%, control 69.1%; z =0.306, p = 0.760, Table 3a). While nestling survival to fledging was higher in prenatally TH supplemented than control nestlings (TH: 83.05% vs CO 69.44%), our model did not provide clear support for such a difference (z = 0.136, p = 0.892, Table 3b) and an analysis from egg to fledgling stage yielded qualitatively similar results (treatment p = 0.86). Fledging success was lower in the nests with larger brood sizes (z = - 2.333, p = 0.020, Table 3b). Juvenile recapture rate was not significantly different across the groups (TH: 28.57% vs CO 34.00%, z=-0.993, p = 0.321, Table 3c).

**Table 3.**
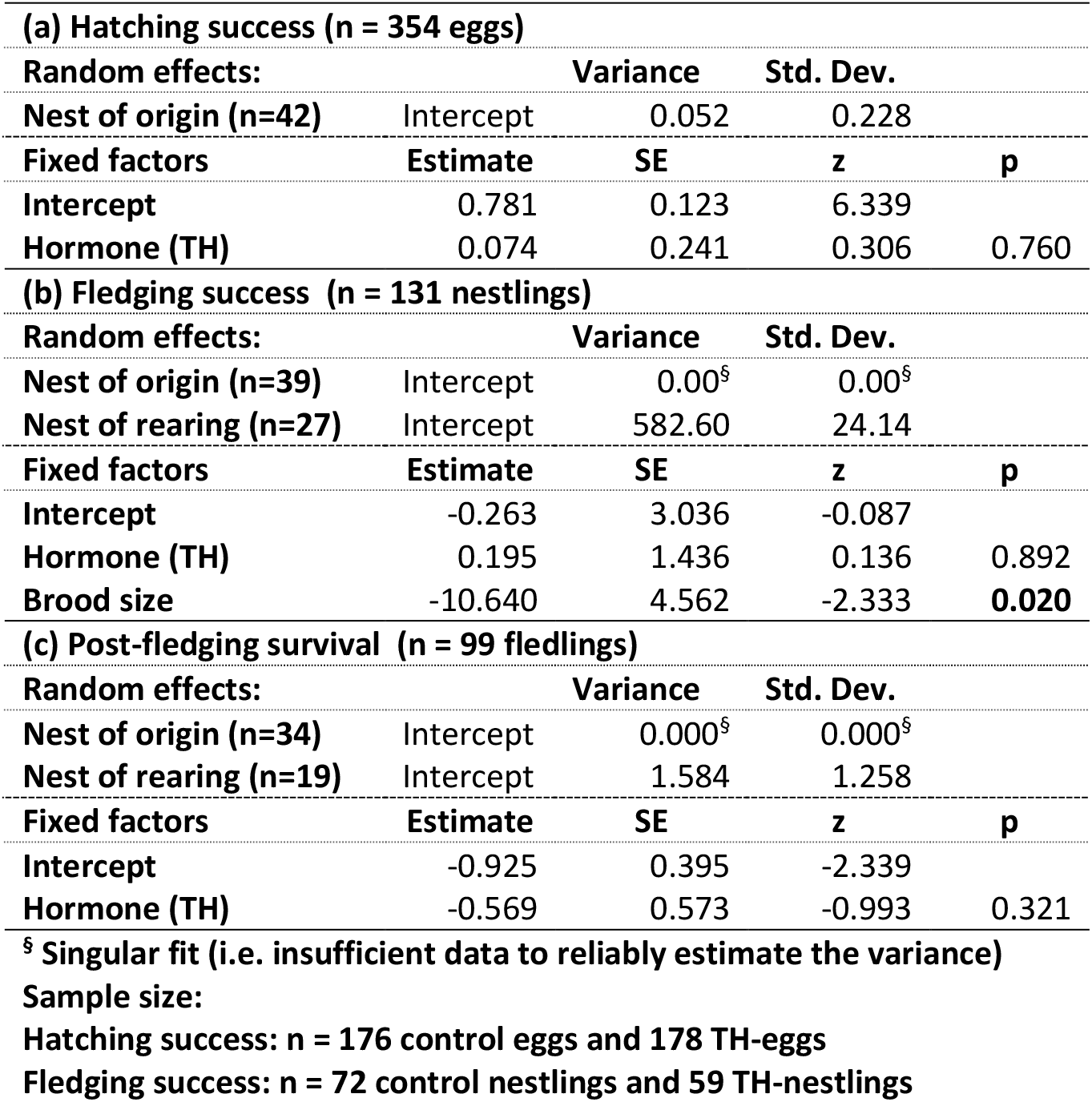
Generalized linear mixed models of hatching success, fledging success, and post-fledging survival. For fledging success and post-fledging survival, only the nestlings that were reared in the nests without temperature manipulation were included. The models were fit by maximum likelihood approach using Laplace approximation with binomial distribution and logit link function. All input variables were mean-centered before model-fitting.

## Discussion

By manipulating prenatal exposure to THs in a wild passerine species, we demonstrate that an increase in prenatal THs can accelerate both postnatal growth and telomere shortening. Yet, we did not detect significant effects of elevated prenatal THs on postnatal oxidative stress levels, cellular energetics measured as mtDNA copy number, circulating TH levels, or short to medium-term survival (*i.e*. hatching success, fledging success, and apparent survival to the next autumn/winter).

Shorter telomeres in the TH-supplemented group were only detected from 14 days after hatching onwards, which seems to exclude direct effects or prenatal TH on telomere dynamics during embryonic development. Yet, this coincides with the accelerated postnatal body mass and tarsus growth observed in the TH-group (between day 7 and 14) compared to controls, and faster growth can accelerate telomere shortening either through enhanced cell division or through inducing oxidative stress (Monaghan and Ozanne, 2018). Considering the lack of impact of prenatal THs on oxidative stress reported here and in previous work on birds (Hsu et al., 2020, 2019a; Sarraude et al., 2020a; Sarraude et al., 2020b), it is not surprising the accelerated telomere shortening observed in this study could not be explained by the measured oxidative stress biomarkers. Yet, we cannot fully rule out this explanation since DNA damage was not directly assessed and oxidative stress sensitivity might vary between different biomolecules (Reichert and Stier, 2017). The effect of prenatal THs on telomere length seemed to increase with age (*i.e*. stronger effect observed in juveniles than at day 14), which could be explained by a delayed effect of fast postnatal growth (day 7 to day 14) because telomeres only shorten at the subsequent cellular division, and therefore delayed effects are likely to be observed (Monaghan and Ozanne, 2018). Nevertheless, including body mass growth between day 7 and day 14 in the model did not explain the difference in telomere shortening. Our hypothesis that prenatal THs could program postnatal metabolic and endocrine function (and thus affect telomere shortening indirectly) was not supported as we found no evidence for differences in mitochondrial density or plasma TH levels (key coordinators of metabolism) across the treatment groups. The latter results are not fully surprising considering the limited evidence supporting a prenatal programming of plasma THs (sex-specific effect on T4 only, Hsu et al. 2017; no effect, Hsu et al. 2020) or mitochondrial density (no effect, Stier et al. 2020; Hsu et al. 2020). Yet, our measurements are limited to blood and tissue-specific effects of TH on mitochondrial density or TH-related cellular machinery (receptors and converting enzymes; Ruuskanen et al. 2022) may occur as this has been shown for instance for glucocorticoids (Weber et al. 2002). According to the *metabolic telomere attrition hypothesis*, telomere shortening is likely to be increased during energy-demanding periods, and accelerated growth under limited resources is likely carrying a metabolic cost (Casagrande and Hau, 2019).

The effects observed here on telomeres in response to increased prenatal TH levels are in sharp contrast with our previous findings in another passerine species, the collared flycatcher (Stier et al. 2020). Indeed, increasing prenatal THs in the collared flycatcher increased telomere length measured very shortly after hatching (day 2), while not affecting telomere shortening during postnatal growth (Stier et al. 2020). There are several alternative explanations for these contrasted findings: (1) we cannot exclude that prenatal THs increased telomere length during embryonic development in great tits since our first telomere length measurement (day 7, *ca*. 70% of fledging body mass) was done considerably later than in collared flycatchers (*i.e*. day 2, *ca*. 20% of fledging body mass). (2) THs differently influenced post-natal growth in the two species: great tits from the TH-supplemented eggs were initially slightly smaller, but grew faster in the late nestling period, whereas collared flycatchers from the TH-supplemented eggs were bigger soon after hatching, but grew slightly slower during the nestling stage (Hsu et al., 2019a). These contrasting growth patterns may explain, at least partly, the contrasted findings in these two species: an increased metabolic demand for fast postnatal growth in great tit in response to prenatal TH could accelerate telomere shortening, while a reduced metabolic demand in collared flycatcher from TH-injected eggs could enable telomere maintenance (Casagrande and Hau, 2019). Measurements of the mTOR signaling pathway could shed light on the validity of this hypothesis (Casagrande and Hau, 2019). (3) The effects of maternal signals on offspring are expected to differ based on environmental conditions, so called context-dependent effects: For example, prenatal androgens have been found to differentially influence offspring growth depending on season or food availability (Groothuis et al., 2020; Muriel et al., 2015). We recently found no evidence for prenatal THs differentially influencing growth and early-life survival depending on rearing temperature (Hsu et al. 2020), yet the influence of resource availability was not tested. In two sister species, collared and pied flycatchers, differential effects of prenatal THs on growth may have been explained by differences in food resources (Sarraude et al. 2020b). (4) These two species exhibit different life-histories, as collared flycatchers are migratory and great tits (relatively) sedentary. Migratory species have generally higher metabolism (Jetz et al., 2008) as well as an overall faster pace of life (Soriano-Redondo et al. 2021), and may thus present differences in TH physiology. We may speculate that the telomere maintenance, and role of THs in the regulation of telomere maintenance may differ across species with different life-histories. Telomere maintenance is known to be influenced by e.g. species lifespan (Haussmann et al., 2004) and migratory populations within a species have shorter telomeres (Bauer et al., 2016). Yet, to our knowledge, studies have not considered migratory vs. non-migratory species in telomere maintenance. Species-specific effects of the prenatal hormonal environment on telomere dynamics have already been described: elevated prenatal glucocorticoid levels led to shorter telomeres in domestic chicken and female zebra finch (Haussmann et al., 2012; Tissier et al., 2014), but to longer telomeres in yellow-legged gull (Noguera, da Silva, and Velando 2020). To understand if any of the hypotheses presented above is likely to explain the contrasted effects found here and in Stier et al. (2020), more studies on the impact of prenatal THs on telomere biology across taxa are needed.

As mentioned above, in the collared flycatcher, elevated yolk THs increased nestlings body mass at day 2 (Hsu et al. 2019), whereas in this great tit study, elevated yolk THs only non-significantly reduced nestling body mass at day 2 (marginal mean ± SE = −0.402 ± 0.245, p=0.109). In both studies, the lighter group of nestlings caught up with the other group before fledging. It therefore appears that the main difference lies on the differential effects of THs during the peri-hatching period. Interestingly, another experiment in the same population of great tits found that elevated yolk THs significantly shortened the time needed to hatch by 0.6 days on average (Cossin-Sevrin et al. 2022). Despite the shorter developmental time, the nestling body mass at day 2 was also found non-significant (Cossin-Sevrin et al. 2022). Thus, the exact peri-hatching effects of prenatal THs in altricial birds still need more information to clarify.

To understand the selective pressures on maternal allocation and signaling, it is important to characterize true fitness effects from development to survival and lifetime reproductive success. In contrast to our predictions, we found no effects of elevated prenatal TH-supplementation on apparent survival or individual condition during their first autumn/winter. Unfortunately, longer-term effects on survival and reproductive success are very challenging to measure in such an experimental setting in the wild, as the recruitment to first breeding is usually very low (e.g. < 10% Radersma et al., 2015), which requires a very high number of nests to be manipulated, something often not feasible for both logistical and ethical reasons. Yet, long-term effects of accelerated early-life telomere shortening may lead to a decrease in longer-term survival and lifespan (Eastwood et al., 2019; Heidinger et al., 2012). The negative effects on later stages suggest that there need to be benefits of high prenatal THs for the offspring or mother, potentially during early-life stages, where selection can be strong. In this study, the elevated growth rates could increase offspring competitive abilities early in life, yet, the effects of prenatal THs on growth seem to be highly inconsistent across studies (Hsu et al., 2020, 2019a, 2017; Ruuskanen et al., 2016a; Sarraude et al., 2020a; Sarraude et al., 2020b). While circulating THs are known to be associated with health and ageing in humans and mammalian models (Bano et al. 2017, 2019; Møllehave et al. 2020), our study is the first to show that exposure to higher prenatal thyroid hormones can lead to accelerated cellular ageing measured through telomere length. This should stimulate further research using both epidemiological and experimental approaches across taxa to uncover the potential regulation of telomere biology by thyroid hormones both pre- and postnatally.

## Author contribution

BYH, AS & SR designed the study and collected field data. NCS, AS & SR conducted laboratory work. BYH analyzed the data. BYH, AS & SR wrote the manuscript, with input from NCS.

## Acknowledgements

This study was financially supported by the Academy of Finland (#286278 to SR). NCS acknowledges support from the EDUFI Fellowship and Maupertuis Grant. B-Y.H work was supported by grants from the Ella and Georg Ehrnrooth Foundation and Academy of Finland (#332716). AS was supported by a ‘Turku Collegium for Science and Medicine’ Fellowship and a Marie Sklodowska-Curie Postdoctoral Fellowship (#894963) at the time of writing. We thank all field assistants, especially Lucas Bousseau, Thomas Rosille, Hsiao-Yin Liu, and Jorma Nurmi for their great effort. All authors declare no conflict of interest.

## Data accessibility

All data is archived and available in Figshare (DOI: 10.6084/m9.figshare.20110409).

---

**Table S1.** Linear mixed models (GLMM) on the effects of prenatal thyroid hormone elevation on nestling body mass and tarsus length at day 2, day 7, and day 14 posthatching. The denominator degree of freedom (ddf) were approximated by the Kenward-Roger method to calculate p values.

| **(a) Body mass at day 2 (n = 218 nestlings)** |  |  |  |  |  |  |
| --- | --- | --- | --- | --- | --- | --- |
| **Random effects:** |  | **Variance** | **Std. Dev.** |  |  |  |
| **Nest of origin (n=41)** | Intercept | 0.481 | 0.694 |  |  |  |
| **Residual** |  | 0.625 | 0.791 |  |  |  |
| **Fixed factors** | **Estimate** | **SE** | **t** | **df** | **F** | **p** |
| **Intercept** | 3.849 | 0.123 | 31.426 |  |  |  |
| **Hormone (TH)** | -0.402 | 0.245 | -1.639 | 1, 38.69 | 2.686 | 0.109 |
| Sample size: n = 99 control and 119 TH-supplemented |  |  |  |  |  |  |
| The total sample size at day 2 (n =218 nestlings from 41 nests) was smaller than that at day 7 because we missed the day 2 for one nest (n = 7 nestlings) |  |  |  |  |  |  |
| **(b) Body mass at day 7 (n = 221 nestlings)** |  |  |  |  |  |  |
| **Random effects:** |  | **Variance** | **Std. Dev.** |  |  |  |
| **Nest of origin (n=41)** | Intercept | 0.944 | 0.972 |  |  |  |
| **Nest of rearing (n=41)** | Intercept | <0.001 | <0.001 |  |  |  |
| **Residual** |  | 1.482 | 1.217 |  |  |  |
| **Fixed factors** | **Estimate** | **SE** | **t** | **df** | **F** | **p** |
| **Intercept** | 12.495 | 0.175 | 71.459 |  |  |  |
| **Hormone (TH)** | -0.367 | 0.353 | -1.040 | 1, 38.93 | 1.081 | 0.305 |
| **Cross-foster (yes)** | -0.347 | 0.179 | -1.945 | 1, 179.22 | 3.782 | 0.053 |
| **Brood size** | -0.141 | 0.072 | -1.956 | 1, 25.71 | 3.828 | 0.061 |
| **Hormone × Cross-foster** | 0.668 | 0.368 | 1.813 | 1, 28.49 | 3.286 | 0.080 |
| **(c) Tarsus length at day 7 (n = 221 nestlings)** |  |  |  |  |  |  |
| **Random effects:** |  | **Variance** | **Std. Dev.** |  |  |  |
| **Nest of origin (n=41)** | Intercept | 0.4916 | 0.700 |  |  |  |
| **Nest of rearing (n=41)** | Intercept | 0.000<sup>§</sup> | 0.000<sup>§</sup> |  |  |  |
| **Residual** |  | 0.982 | 0.991 |  |  |  |
| **Fixed factors** | **Estimate** | **SE** | **t** | **df** | **F** | **p** |
| **Intercept** | 18.940 | 0.205 | 92.592 |  |  |  |
| **Hormone (TH)** | -0.275 | 0.322 | -0.855 | 1, 42.78 | 0.731 | 0.397 |
| **Cross-foster (yes)** | -0.330 | 0.146 | -2.268 | 1, 178.84 | 5.145 | 0.025 |
| **Proportion of TH nestlings** | -0.297 | 0.482 | -0.617 | 1, 54.90 | 0.381 | 0.540 |
| **Measurer 2** | 0.211 | 0.199 | 1.059 | 2, 204.00 | 3.357 | 0.037 |
| **Measurer 3** | 0.523 | 0.204 | 2.561 |  |  |  |
| **Hormone × proportion of TH nestlings** | -1.711 | 0.972 | -1.762 | 1, 58.65 | 3.103 | 0.083 |

Table S1 (cont.)
| **(d) Body mass at day 14 (n = 99 nestlings)** |  |  |  |  |  |  |
| --- | --- | --- | --- | --- | --- | --- |
| **Random effects:** |  | **Variance** | **Std. Dev.** |  |  |  |
| **Nest of origin (n=34)** | Intercept | 0.310 | 0.557 |  |  |  |
| **Nest of rearing (n=19)** | Intercept | 0.228 | 0.477 |  |  |  |
| **Residual** |  | 0.801 | 0.895 |  |  |  |
| **Fixed factors** | **Estimate** | **SE** | **t** | **df** | **F** | **p** |
| **Intercept** | 18.603 | 0.176 | 105.693 |  |  |  |
| **Hormone (TH)** | 0.099 | 0.303 | 0.747 | 1, 22.45 | 0.130 | 0.722 |
| **(e) Tarsus length at day 14 (n = 99 nestlings)** |  |  |  |  |  |  |
| **Random effects:** |  | **Variance** | **Std. Dev.** |  |  |  |
| **Nest of origin (n=34)** | Intercept | 0.132 | 0.363 |  |  |  |
| **Nest of rearing (n=19)** | Intercept | 0.000<sup>§</sup> | 0.000<sup>§</sup> |  |  |  |
| **Residual** |  | 0.283 | 0.532 |  |  |  |
| **Fixed factors** | **Estimate** | **SE** | **t** | **df** | **F** | **p** |
| **Intercept** | 23.199 | 0.238 | 97.313 |  |  |  |
| **Hormone (TH)** | 0.157 | 0.175 | 0.899 | 1, 27.77 | 0.808 | 0.376 |
| **Measurer 2** | -0.488 | 0.265 | -1.845 | 2, 61.90 | 4.014 | 0.023 |
| **Measurer 3** | -0.066 | 0.256 | -0.258 |  |  |  |

Sample sizes: n = 50 control and 49 TH-supplemented
<sup>§</sup> Singular fit (i.e. insufficient data to reliably estimate the variance)

**Table S2.** Linear mixed models of blood oxidative stress biomarkers in response to prenatal TH-supplementation.

| **(a) total GSH $\mu\text{mol}/\text{mg}$ protein (n = 58 nestlings)** |  |  |  |  |  |  |
| --- | --- | --- | --- | --- | --- | --- |
| **Random effects:** |  | **Variance** | **Std. Dev.** |  |  |  |
| **Nest of origin (n=29)** | Intercept | 0.031 | 0.177 |  |  |  |
| **Nest of rearing (n=18)** | Intercept | 0.050 | 0.223 |  |  |  |
| **Residual** |  | 0.044 | 0.210 |  |  |  |
| **Fixed factors** | **Estimate** | **SE** | **t** | **df** | **F** | **p** |
| **Intercept** | -1.492 | 0.114 | -13.148 |  |  |  |
| **Hormone (TH)** | 0.014 | 0.105 | 0.128 | 1, 15.46 | 0.016 | 0.900 |
| **GSH assay batch** | 0.221 | 0.145 | 1.526 | 1, 16.57 | 2.328 | 0.146 |
| **(b) GSH:GSSG ratio (n = 58 nestlings)** |  |  |  |  |  |  |
| **Random effects:** |  | **Variance** | **Std. Dev.** |  |  |  |
| **Nest of origin (n=29)** | Intercept | 0.210 | 0.458 |  |  |  |
| **Nest of rearing (n=18)** | Intercept | 0.031 | 0.175 |  |  |  |
| **Residual** |  | 0.134 | 0.367 |  |  |  |
| **Fixed factors** | **Estimate** | **SE** | **t** | **df** | **F** | **p** |
| **Intercept** | -3.106 | 0.181 | -17.207 |  |  |  |
| **Hormone (TH)** | -0.171 | 0.243 | -0.703 | 1, 16.75 | 0.456 | 0.509 |
| **Body mass** | 0.106 | 0.059 | 1.794 | 1, 41.88 | 2.986 | 0.091 |
| **Proportion of TH-nestlings** | -0.173 | 0.462 | -0.375 | 1, 21.30 | 0.124 | 0.728 |
| **GSH assay batch** | 0.133 | 0.237 | 0.561 | 1, 16.87 | 0.307 | 0.587 |
| **Hormone × Proportion of TH-nestlings** | 1.787 | 0.926 | 1.930 | 1, 19.81 | 3.553 | 0.074 |
| **(c) MDA $\text{nmol}/\mu\text{g}$ protein (n = 59 nestlings)** |  |  |  |  |  |  |
| **Random effects:** |  | **Variance** | **Std. Dev.** |  |  |  |
| **Nest of origin (n=30)** | Intercept | 0.004 | 0.059 |  |  |  |
| **Nest of rearing (n=18)** | Intercept | 0.004 | 0.060 |  |  |  |
| **TBARS batch (n = 6)** | Intercept | 0.100 | 0.317 |  |  |  |
| **Residual** |  | 0.039 | 0.198 |  |  |  |
| **Fixed factors** | **Estimate** | **SE** | **t** | **df** | **F** | **p** |
| **Intercept** | -3.286 | 0.135 | -24.340 |  |  |  |
| **Hormone (TH)** | 0.036 | 0.067 | 0.536 | 1, 16.86 | 0.287 | 0.599 |

Sample sizes:
total GSH and GSH:GSSG ratio, n = 26 control and 32 TH-supplemented
MDA, n = 26 control and 33 TH-supplemented

**Table S3.** Linear mixed models of blood plasma thyroid hormones growth in response to prenatal TH-supplementation.

| **(a) T3 pmol/ml (n = 24 nestlings)** |  |  |  |  |  |  |
| --- | --- | --- | --- | --- | --- | --- |
| **Random effects:** |  | **Variance** | **Std. Dev.** |  |  |  |
| **Nest of origin (n=18)** | Intercept | 0.054 | 0.232 |  |  |  |
| **Nest of rearing (n=11)** | Intercept | 0.127 | 0.357 |  |  |  |
| **Residual** |  | 0.132 | 0.363 |  |  |  |
| **Fixed factors** | **Estimate** | **SE** | **t** | **df** | **F** | **p** |
| **Intercept** | 1.218 | 0.147 | 8.273 |  |  |  |
| **Hormone (TH)** | -0.026 | 0.202 | -0.128 | 1, 7.37 | 0.016 | 0.902 |
| **(b) T4 pmol/ml (n = 24 nestlings)** |  |  |  |  |  |  |
| **Random effects:** |  | **Variance** | **Std. Dev.** |  |  |  |
| **Nest of origin (n=18)** | Intercept | 0.738 | 0.859 |  |  |  |
| **Nest of rearing (n=11)** | Intercept | 2.934 | 1.713 |  |  |  |
| **Residual** |  | 5.721 | 2.392 |  |  |  |
| **Fixed factors** | **Estimate** | **SE** | **t** | **df** | **F** | **p** |
| **Intercept** | 7.452 | 0.758 | 9.837 |  |  |  |
| **Hormone (TH)** | -0.041 | 1.113 | -0.037 | 1, 6.57 | 0.001 | 0.973 |
| **Sex** | 2.101 | 1.119 | 1.877 | 1, 14.99 | 2.532 | 0.132 |

Sample sizes: n = 11 control and 13 TH-supplemented

## References

Aljanabi, S.M., Martinez, I., 1997. Universal and rapid salt-extraction of high quality genomic DNA for PCR-based techniques. Nucleic Acids Research. https://doi.org/10.1093/nar/25.22.4692

Arbeev, K.G., Verhulst, S., Steenstrup, T., Kark, J.D., Bagley, O., Kooperberg, C., Reiner, A.P., Hwang, S.-J., Levy, D., Fitzpatrick, A.L., Christensen, K., Yashin, A.I., Aviv, A., 2020. Association of Leukocyte Telomere Length With Mortality Among Adult Participants in 3 Longitudinal Studies. JAMA Netw Open 3, e200023. https://doi.org/10.1001/jamanetworkopen.2020.0023

Atema, E., Mulder, E., van Noordwijk, A.J., Verhulst, S., 2019. Ultralong telomeres shorten with age in nestling great tits but are static in adults and mask attrition of short telomeres. Mol Ecol Resour 19, 648–658. https://doi.org/10.1111/1755-0998.12996

Bano, A., Chaker, L., Mattace-Raso, F.U.S., Terzikhan, N., Kavousi, M., Ikram, M.A., Peeters, R.P., Franco, O.H., 2019. Thyroid function and life expectancy with and without noncommunicable diseases: A population-based study. PLoS Med 16, e1002957. https://doi.org/10.1371/journal.pmed.1002957

Bano, A., Dhana, K., Chaker, L., Kavousi, M., Ikram, M.A., Mattace-Raso, F.U.S., Peeters, R.P., Franco, O.H., 2017. zeta Association of Thyroid Function With Life Expectancy With and Without Cardiovascular Disease The Rotterdam Study. Jama Internal Medicine. https://doi.org/10.1001/jamainternmed.2017.4836

Barnes, S.K., Ozanne, S.E., 2011. Pathways linking the early environment to long-term health and lifespan. Progress in Biophysics and Molecular Biology 106, 323–336. https://doi.org/10.1016/j.pbiomolbio.2010.12.005

Bates, D., Machler, M., Bolker, B.M., Walker, S.C., 2015. Fitting Linear Mixed-Effects Models Using lme4. Journal of Statistical Software. https://doi.org/10.18637/jss.v067.i01

Bauer, C.M., Heidinger, B.J., Ketterson, E.D., Greives, T.J., 2016. A migratory lifestyle is associated with shorter telomeres in a songbird (Junco hyemalis). Auk. https://doi.org/10.1642/auk-16-56.1

Casagrande, S., Hau, M., 2019. Telomere attrition: metabolic regulation and signalling function? Biology Letters. https://doi.org/10.1098/rsbl.2018.0885

Chang, H.W., Cheng, C.A., Gu, D.L., Chang, C.C., Su, S.H., Wen, C.H., Chou, Y.C., Chou, T.C., Yao, C.T., Tsai, C.L., Cheng, C.C., 2008. High-throughput avian molecular sexing by SYBR green-based real-time PCR combined with melting curve analysis. Bmc Biotechnology. https://doi.org/10.1186/1472-6750-8-12

Cioffi, F., Senese, R., Lanni, A., Goglia, F., 2013. Thyroid hormones and mitochondria: With a brief look at derivatives and analogues. Molecular and Cellular Endocrinology. https://doi.org/10.1016/j.mce.2013.06.006

Cossin-Sevrin, N., Hsu, B.-Y., Marciau, C., Viblanc, V.A., Ruuskanen, S., Stier, A., 2022. Effect of prenatal glucocorticoids and thyroid hormones on developmental plasticity of mitochondrial aerobic metabolism, growth and survival: an experimental test in wild great tits. Journal of Experimental Biology 225, jeb243414. https://doi.org/10.1242/jeb.243414

Criscuolo, F., Torres, R., Zahn, S., Williams, T.D., 2020. Telomere dynamics from hatching to sexual maturity and maternal effects in the ‘multivariate egg.’ Journal of Experimental Biology 223, jeb232496. https://doi.org/10.1242/jeb.232496

Eastwood, J.R., Hall, M.L., Teunissen, N., Kingma, S.A., Aranzamendi, N.H., Fan, M., Roast, M., Verhulst, S., Peters, A., 2019. Early-life telomere length predicts lifespan and lifetime reproductive success in a wild bird. Molecular Ecology. https://doi.org/10.1111/mec.15002

Ellegren, H., Fridolfsson, A.K., 1997. Male-driven evolution of DNA sequences in birds. Nat Genet 17, 182–184. https://doi.org/10.1038/ng1097-182

Entringer, S., de Punder, K., Buss, C., Wadhwa, P.D., 2018. The fetal programming of telomere biology hypothesis: an update. Philos Trans R Soc Lond B Biol Sci 373, 20170151. https://doi.org/10.1098/rstb.2017.0151

Espin, S., Sanchez Virosta, P., García-Fernández, A., Eeva, T., 2017. A microplate adaptation of the thiobarbituric acid reactive substances assay to determine lipid peroxidation fluorometrically in small sample volumes. Revista de Toxicologia 34, In press.

Godfrey, K.M., Barker, D.J., 2001. Fetal programming and adult health. Public Health Nutr 4, 611–624. https://doi.org/10.1079/phn2001145

Groothuis, T.G.G., Kumar, N., Hsu, B.Y., 2020. Explaining discrepancies in the study of maternal effects: the role of context and embryo. Current Opinion in Behavioral Sciences. https://doi.org/10.1016/j.cobeha.2020.10.006

Halekoh, U., Hojsgaard, S., 2014. Kenward-Roger Approximation and Parametric Bootstrap Methods for Tests in Linear Mixed Models - The R Package pbkrtest. Journal of Statistical Software.

Hartig, F. (2021). DHARMa: Residual Diagnostics for Hierarchical (Multi-Level / Mixed) Regression Models. R package version 0.4.3. https://CRAN.R-project.org/package=DHARMa

Haussmann, M.F., Longenecker, A.S., Marchetto, N.M., Juliano, S.A., Bowden, R.M., 2012. Embryonic exposure to corticosterone modifies the juvenile stress response, oxidative stress and telomere length. Proceedings of the Royal Society B-Biological Sciences. https://doi.org/10.1098/rspb.2011.1913

Haussmann, M.F., Winkler, D.W., Huntington, C.E., Nisbet, I.C.T., Vleck, C.M., 2004. Telomerase expression is differentially regulated in birds of differing life span. Strategies for Engineered Negligible Senescence: Why Genuine Control of Aging May Be Foreseeable. https://doi.org/10.1196/annals.1297.029

Heidinger, B.J., Blount, J.D., Boner, W., Griffiths, K., Metcalfe, N.B., Monaghan, P., 2012. Telomere length in early life predicts lifespan. Proceedings of the National Academy of Sciences 109, 1743–1748. https://doi.org/10.1073/pnas.1113306109

Hsu, B.Y., Dijkstra, C., Darras, V.M., de Vries, B., Groothuis, T.G.G., 2017. Maternal thyroid hormones enhance hatching success but decrease nestling body mass in the rock pigeon (Columba livia). General and Comparative Endocrinology. https://doi.org/10.1016/j.ygcen.2016.10.011

Hsu, B.Y., Doligez, B., Gustafsson, L., Ruuskanen, S., 2019a. Transient growth-enhancing effects of elevated maternal thyroid hormones at no apparent oxidative cost during early postnatal period. Journal of Avian Biology. https://doi.org/10.1111/jav.01919

Hsu, B.Y., Sarraude, T., Cossin-Sevrin, N., Crombecque, M., Stier, A., Ruuskanen, S., 2020. Testing for context-dependent effects of prenatal thyroid hormones on offspring survival and physiology: an experimental temperature manipulation. Scientific Reports. https://doi.org/10.1038/s41598-020-71511-y

Hsu, B.Y., Verhagen, I., Gienapp, P., Darras, V.M., Visser, M.E., Ruuskanen, S., 2019b. Between-and Within-Individual Variation of Maternal Thyroid Hormone Deposition in Wild Great Tits (Parus major). American Naturalist. https://doi.org/10.1086/704738

Jetz, W., Freckleton, R.P., McKechnie, A.E., 2008. Environment, Migratory Tendency, Phylogeny and Basal Metabolic Rate in Birds. Plos One. https://doi.org/10.1371/journal.pone.0003261

Kärkkäinen, T., Briga, M., Laaksonen, T., Stier, A., n.d. Within-individual repeatability in telomere length: A meta-analysis in nonmammalian vertebrates. Molecular Ecology n/a. https://doi.org/10.1111/mec.16155

Kuznetsova, A., Brockhoff, P.B., Christensen, R.H.B., 2017. lmerTest Package: Tests in Linear Mixed Effects Models. Journal of Statistical Software 82, 1–26. https://doi.org/10.18637/jss.v082.i13

Larsen, S., Nielsen, J., Hansen, C.N., Nielsen, L.B., Wibrand, F., Stride, N., Schroder, H.D., Boushel, R., Helge, J.W., Dela, F., Hey-Mogensen, M., 2012. Biomarkers of mitochondrial content in skeletal muscle of healthy young human subjects. J Physiol 590, 3349–3360. https://doi.org/10.1113/jphysiol.2012.230185

Length, R.V. (2021). emmeans: Estimated Marginal Means, aka Least-Squares Means. R package version 1.6.2-1. https://CRAN.R-project.org/package=emmeans

Liu, J.S., Chen, Y.Q., Li, M., 2006. Thyroid hormones increase liver and muscle thermogenic capacity in the little buntings (Emberiza pusilla). Journal of Thermal Biology. https://doi.org/10.1016/j.jtherbio.2006.01.002

López-Otín, C., Blasco, M.A., Partridge, L., Serrano, M., Kroemer, G., 2013. The Hallmarks of Aging. Cell 153, 1194–1217. https://doi.org/10.1016/j.cell.2013.05.039

Marchetto, N.M., Glynn, R.A., Ferry, M.L., Ostojic, M., Wolff, S.M., Yao, R., Haussmann, M.F., 2016. Prenatal stress and newborn telomere length. American Journal of Obstetrics and Gynecology 215, 94.e1–94.e8. https://doi.org/10.1016/j.ajog.2016.01.177

McNabb, F.M.A., 2007. The hypothalamic-pituitary-thyroid (HPT) axis in birds and its role in bird development and reproduction. Critical Reviews in Toxicology. https://doi.org/10.1080/10408440601123552

Medici, M., Timmermans, S., Visser, W., Keizer-Schrama, S., Jaddoe, V.W.W., Hofman, A., Hooijkaas, H., de Rijke, Y.B., Tiemeier, H., Bongers-Schokking, J.J., Visser, T.J., Peeters, R.P., Steegers, E.A.P., 2013. Maternal Thyroid Hormone Parameters during Early Pregnancy and Birth Weight: The Generation R Study. Journal of Clinical Endocrinology & Metabolism. https://doi.org/10.1210/jc.2012-2420

Metcalfe, N.B., Monaghan, P., 2001. Compensation for a bad start: grow now, pay later? Trends in Ecology & Evolution.

Møllehave, L.T., Skaaby, T., Linneberg, A., Knudsen, N., Jørgensen, T., Thuesen, B.H., 2020. The association of thyroid stimulation hormone levels with incident ischemic heart disease, incident stroke, and all-cause mortality. Endocrine 68, 358–367. https://doi.org/10.1007/s12020-020-02216-5

Monaghan, P., Ozanne, S.E., 2018. Somatic growth and telomere dynamics in vertebrates: relationships, mechanisms and consequences. Philos Trans R Soc Lond B Biol Sci 373, 20160446. https://doi.org/10.1098/rstb.2016.0446

Mullur, R., Liu, Y.-Y., Brent, G.A., 2014. Thyroid Hormone Regulation of Metabolism. Physiological Reviews 94, 355–382. https://doi.org/10.1152/physrev.00030.2013

Muñoz-Lorente, M.A., Cano-Martin, A.C., Blasco, M.A., 2019. Mice with hyper-long telomeres show less metabolic aging and longer lifespans. Nat Commun 10, 4723. https://doi.org/10.1038/s41467-019-12664-x

Muriel, J., Salmón, P., Nunez-Buiza, A., de Salas, F., Perez-Rodriguez, L., Puerta, M., Gil, D., 2015. Context-dependent effects of yolk androgens on nestling growth and immune function in a multibrooded passerine. Journal of Evolutionary Biology. https://doi.org/10.1111/jeb.12668

Noguera, J.C., da Silva, A., Velando, A., n.d. Egg corticosterone can stimulate telomerase activity and promote longer telomeres during embryo development. Molecular Ecology n/a. https://doi.org/10.1111/mec.15694

Parolini, M., Possenti, C.D., Caprioli, M., Rubolini, D., Romano, A., Saino, N., 2019. Egg Testosterone Differentially Affects Telomere Length in Somatic Tissues of Yellow-Legged Gull Embryos. Physiological and Biochemical Zoology. https://doi.org/10.1086/705037

Podmokla, E., Drobniak, S.M., Rutkowska, J., 2018. Chicken or egg? Outcomes of experimental manipulations of maternally transmitted hormones depend on administration method - a meta-analysis. Biological Reviews. https://doi.org/10.1111/brv.12406

Radersma, R., Komdeur, J., Tinbergen, J.M., 2015. Early morning fledging improves recruitment in Great Tits Parus major. Ibis 157, 351–355. https://doi.org/10.1111/ibi.12230

Reichert, S., Stier, A., 2017. Does oxidative stress shorten telomeres in vivo? A review. Biology Letters. https://doi.org/10.1098/rsbl.2017.0463

Ruuskanen, S., Darras, V.M., Visser, M.E., Groothuis, T.G.G., 2016a. Effects of experimentally manipulated yolk thyroid hormone levels on offspring development in a wild bird species. Hormones and Behavior. https://doi.org/10.1016/j.yhbeh.2016.03.006

Ruuskanen, S., Espin, S., Sanchez-Virosta, P., Sarraude, T., Hsu, B.Y., Pajunen, P., Costa, R.A., Eens, M., Hargitai, R., Torok, J., Eeva, T., 2019. Transgenerational endocrine disruption: Does elemental pollution affect egg or nestling thyroid hormone levels in a wild songbird? Environmental Pollution. https://doi.org/10.1016/j.envpol.2019.01.088

Ruuskanen, S., Groothuis, T.G.G., Schaper, S.V., Darras, V.M., de Vries, B., Visser, M.E., 2016b. Temperature-induced variation in yolk androgen and thyroid hormone levels in avian eggs. General and Comparative Endocrinology. https://doi.org/10.1016/j.ygcen.2016.05.026

Ruuskanen, S., Hsu, B.Y., 2018. Maternal Thyroid Hormones: An Unexplored Mechanism Underlying Maternal Effects in an Ecological Framework. Physiological and Biochemical Zoology. https://doi.org/10.1086/697380

Ruuskanen, S., Hsu, B.-Y., Heinonen, A., Darras, V., Vainio, M., Sarraude, T., Rokka, A., 2018. A new method for measuring thyroid hormones using nano-LC-MS/MS. Journal of Chromatography B.

Salmón, P., Nilsson, J.F., Nord, A., Bensch, S., Isaksson, C., 2016. Urban environment shortens telomere length in nestling great tits, Parus major. Biol Lett 12, 20160155. https://doi.org/10.1098/rsbl.2016.0155

Salmón, P., Nilsson, J.F., Watson, H., Bensch, S., Isaksson, C., 2017. Selective disappearance of great tits with short telomeres in urban areas. Proc Biol Sci 284, 20171349. https://doi.org/10.1098/rspb.2017.1349

Sarraude, T., Hsu, B.-Y., Groothuis, T., Ruuskanen, S., 2020a. Manipulation of prenatal thyroid hormones does not influence growth or physiology in nestling pied flycatchers. Physiological and Biochemical Zoology.

Sarraude, T., Hsu, B.Y., Groothuis, T., Ruuskanen, S., 2020b. Testing the short-and long-term effects of elevated prenatal exposure to different forms of thyroid hormones. Peerj. https://doi.org/10.7717/peerj.10175

Schielzeth, H., 2010. Simple means to improve the interpretability of regression coefficients. Methods in Ecology and Evolution 1, 103–113. https://doi.org/10.1111/j.2041-210X.2010.00012.x

Schielzeth, H., Dingemanse, N.J., Nakagawa, S., Westneat, D.F., Allegue, H., Teplitsky, C., Réale, D., Dochtermann, N.A., Garamszegi, L.Z., Araya-Ajoy, Y.G., 2020. Robustness of linear mixed-effects models to violations of distributional assumptions. Methods in Ecology and Evolution 11, 1141–1152. https://doi.org/10.1111/2041-210X.13434

Smith, S.M., Nager, R.G., Costantini, D., 2016. Meta-analysis indicates that oxidative stress is both a constraint on and a cost of growth. Ecol Evol 6, 2833–2842. https://doi.org/10.1002/ece3.2080

Stier, A., Bize, P., Hsu, B.Y., Ruuskanen, S., 2019. Plastic but repeatable: rapid adjustments of mitochondrial function and density during reproduction in a wild bird species. Biology Letters. https://doi.org/10.1098/rsbl.2019.0536

Stier, A., Delestrade, A., Bize, P., Zahn, S., Criscuolo, F., Massemin, S., 2016. Investigating how telomere dynamics, growth and life history covary along an elevation gradient in two passerine species. Journal of Avian Biology 47, 134–140. https://doi.org/10.1111/jav.00714

Stier, A., Hsu, B.Y., Marciau, C., Doligez, B., Gustafsson, L., Bize, P., Ruuskanen, S., 2020. Born to be young? Prenatal thyroid hormones increase early-life telomere length in wild collared flycatchers. Biology Letters. https://doi.org/10.1098/rsbl.2020.0364

Stier, A., Massemin, S., Criscuolo, F., 2014. Chronic mitochondrial uncoupling treatment prevents acute cold-induced oxidative stress in birds. Journal of Comparative Physiology B-Biochemical Systems and Environmental Physiology. https://doi.org/10.1007/s00360-014-0856-6

Stier, A., Massemin, S., Zahn, S., Tissier, M.L., Criscuolo, F., 2015. Starting with a handicap: effects of asynchronous hatching on growth rate, oxidative stress and telomere dynamics in free-living great tits. Oecologia. https://doi.org/10.1007/s00442-015-3429-9

Stier, A., Metcalfe, N.B., Monaghan, P., 2020. Pace and stability of embryonic development affect telomere dynamics: an experimental study in a precocial bird model. Proc Biol Sci 287, 20201378. https://doi.org/10.1098/rspb.2020.1378

Tissier, M.L., Williams, T.D., Criscuolo, F., 2014. Maternal Effects Underlie Ageing Costs of Growth in the Zebra Finch (Taeniopygia guttata). PLoS One 9, e97705. https://doi.org/10.1371/journal.pone.0097705

Tschirren, B., Saladin, V., Fitze, P.S., Schwabl, H., Richner, H., 2005. Maternal yolk testosterone does not modulate parasite susceptibility or immune function in great tit nestlings. Journal of Animal Ecology 74: 675–682.

Verhulst, S., 2020. Improving comparability between qPCR-based telomere studies. Molecular Ecology Resources 20, 11–13. https://doi.org/10.1111/1755-0998.13114

Vrijkotte, T.G.M., Hrudey, E.J., Twickler, M.B., 2017. Early Maternal Thyroid Function During Gestation Is Associated With Fetal Growth, Particularly in Male Newborns. The Journal of Clinical Endocrinology & Metabolism 102, 1059–1066. https://doi.org/10.1210/jc.2016-3452

Weber, K., Brück, P., Mikes, Z., Küpper, J-H., Klingenspor, M., Wiesner, R.J.. 2002. Glucocorticoid Hormone Stimulates Mitochondrial Biogenesis Specifically in Skeletal Muscle. Endocrinology, 143, 177–184

Welcker, J., Chastel, O., Gabrielsen, G.W., Guillaumin, J., Kitaysky, A.S., Speakman, J.R., Tremblay, Y., Bech, C., 2013. Thyroid Hormones Correlate with Basal Metabolic Rate but Not Field Metabolic Rate in a Wild Bird Species. Plos One. https://doi.org/10.1371/journal.pone.0056229

Wilbourn, R.V., Moatt, J.P., Froy, H., Walling, C.A., Nussey, D.H., Boonekamp, J.J., 2018. The relationship between telomere length and mortality risk in non-model vertebrate systems: a meta-analysis. Philosophical Transactions of the Royal Society B-Biological Sciences. https://doi.org/10.1098/rstb.2016.0447

Zhang, C., Yang, X., Zhang, Y., Guo, F., Yang, S., Peeters, R.P., Korevaar, T.I.M., Fan, J., Huang, H.-F., 2019. Association Between Maternal Thyroid Hormones and Birth Weight at Early and Late Pregnancy. The Journal of Clinical Endocrinology & Metabolism 104, 5853–5863. https://doi.org/10.1210/jc.2019-00390

